# Nucleolar stress in *Drosophila* neuroblasts, a model for human ribosomopathies

**DOI:** 10.1101/704734

**Authors:** Sonu Shrestha Baral, Molly E. Lieux, Patrick J. DiMario

## Abstract

Different stem cells or progenitor cells display variable threshold requirements for functional ribosomes. For instance, select embryonic neural crest cells or adult bone marrow stem cells, but not others, show lethality due to failures in ribosome biogenesis or function (known as nucleolar stress) in several human ribosomopathies. To determine if various *Drosophila* neuroblasts display differential sensitivities to nucleolar stress, we used CRISPR-Cas9 to disrupt the *Nopp140* gene that encodes two ribosome biogenesis factors (RBFs). Disruption of *Nopp140* induced nucleolar stress that arrested larvae in the second instar stage. While the majority of larval neuroblasts arrested development, the Mushroom Body (MB) neuroblasts continued to proliferate as shown by their maintenance of deadpan, a neuroblast-specific transcription factor, and by their continued EdU incorporation. MB neuroblasts in wild type larvae contained more fibrillarin and Nopp140 in their nucleoli as compared to other neuroblasts, indicating that MB neuroblasts stockpile RBFs as they proliferate in late embryogenesis while other neuroblasts normally enter quiescence. A greater abundance of Nopp140 encoded by maternal transcripts in *Nopp140−/−* MB neuroblasts likely rendered these cells more resilient to nucleolar stress.

**Summary Statement:** Nucleolar stress (loss of ribosome production/function) in certain human stem cells or progenitor cells results in disease. In fruit flies, larval Mushroom Body neuroblasts are relatively resilient to nucleolar stress.

## INTRODUCTION

The nucleolus is the nuclear sub-compartment responsible for ribosomal subunit biogenesis (Baßler and Hurt, 2019). Functional ribosomes in the cytoplasm of eukaryotic cells consist of the small ribosomal subunit with its 18S ribosomal RNA (rRNA) assembled with 33 ribosomal proteins and the large ribosomal subunit with its 28S, 5.8S, and 5S rRNAs assembled with 47 ribosomal proteins. This assembly is a complex choreography of reactions and interactions that begins with RNA polymerase I (RNA Pol I) as it synthesizes pre-rRNA from tandemly repeated ribosomal DNA (rDNA) genes. The 38S pre-rRNA in *Drosophila* undergoes endonuclease cleavages to generate 18S, 5.8S+2S, and 28S rRNAs (Long and Dawid, 1980). These rRNAs are chemically modified by box C/D small nucleolar ribonucleoprotein complexes (snoRNPs) (2’-*O*-methylation) and box H/ACA snoRNPs (pseudouridylation) (Wang et al., 2002; Yang et al., 2000; Bachellerie et al., 2002). Besides endonucleases and snoRNPs, subunit biogenesis requires a myriad of other factors serving as RNP chaperones, RNA helicases, and GTPase release factors (Kressler et al., 2010). The chaperones, often referred to as ribosome biogenesis factors (RBFs), act early in ribosome assembly; they include Nopp140 (Nucleolar and Cajal body phosphoprotein of 140 kDa) and treacle. While both Nopp140 and treacle are found in vertebrates, only Nopp140 orthologues are expressed in all eukaryotes.

Ribosome biogenesis requires high energy expenditures by the cell; approximately 60% of total cellular transcription is devoted to rRNA, with some 2000 ribosomes assembled per minute in actively growing yeast cells (Warner 1999; Woolford and Baserga, 2013). Any perturbation in ribosome biogenesis disrupts cell homeostasis; this is now called nucleolar (or ribosome) stress (Golomb et al., 2014; Tsai and Pederson, 2014; Yang et al., 2018). In humans, nucleolar stress due to mutations in ribosome biogenesis factors (RBFs), processing snoRNPs, or the ribosomal proteins themselves results in disease states collectively called ribosomopathies, of which there are several (Narla and Ebert, 2010). While each ribosomopathy has its own distinct phenotypes, and several display tissue-specificity (McCann and Baserga, 2013), there are commonalities among them: the most prevalent dysfunctions include craniofacial abnormalities, other skeletal defects, and bone marrow failures. All ribosomopathies affect only certain stem cells or progenitors despite the mutation being systemic.

One of these ribosomopathies is the Treacher Collins Syndrome (TCS), a congenital birth defect caused by haplo-insufficiency mutations in the *TCOF1* gene that encodes treacle (Sakai and Trainor, 2009). A particular set of neural crest cells that normally migrate to and populate pharyngeal arches I and II on day 24-25 human embryogenesis have insufficient functional ribosomes in TCS individuals. This leads to p53-dependent apoptosis (Jones et al., 2008). Loss of these particular neural crest cells causes the craniofacial defects. A TCS-like phenotype can also result from mutations in genes encoding RNA Pol I and III subunit proteins, POLR1D and POLR1C respectively (Dauwerse et al., 2011; Noack Watt et al., 2016). The question is, why are only certain progenitor cells affected while others remain resilient?

To investigate the underlying mechanism contributing to stem cell or progenitor cell specificity as seen in the human ribosomopathies, we initiated a study of nucleolar stress in *Drosophila* larval neuroblasts. We wanted to determine if all neuroblast types respond similarly or differentially to nucleolar stress. We typically induce nucleolar stress by depleting Nopp140. Like treacle, metazoan Nopp140 orthologues contain alternating acidic and basic motifs constituting a large central domain of low sequence complexity (Meier, 1996). Treacle and Nopp140 also share similar roles in chaperoning C/D-box snoRNPs to the dense fibrillar component of nucleoli where pre-rRNA is modified by site-specific 2’-*O*-methylation. Unlike treacle, Nopp140 locates to Cajal bodies; thus Nopp140 may also play a role in snoRNP assembly and transport to nucleoli (Gonzales et al., 2005; Hayano et al., 2003; He et al., 2015). With crucial roles in ribosome biogenesis, Nopp140 depletion in *Drosophila* induces nucleolar stress such that cell death occurs either by apoptosis in progenitor imaginal disc cells or by autophagy in terminally differentiated polyploid gut cells (James et al., 2013, 2014).

The *Drosophila* larval brain comprises a diverse set of distinctive neuroblast (NB) lineages generated from a fixed set of founder NBs (Homem and Knoblich, 2012; Hartenstein and Wodarz, 2013). Briefly, there are four major neuroblast types in the *Drosophila* larval brain; Type I NBs, Type II NBs, Mushroom Body (MB) NBs, and Optic Lobe NBs (Fig. 1A). We hypothesize that upon nucleolar stress caused by the loss of Nopp140, different neuroblast lineages exhibit variable phenotypes ranging from a mild loss of lineage progeny cells to substantial loss of the lineage altogether. Here we show that MB neuroblasts are more resilient to the effects of nucleolar stress compared to other neuroblast types. Hence, different neuroblast lineages respond variably to nucleolar stress which is reminiscent of the neural crest cell-specific effects caused by the loss of treacle in TCS individuals.

**Fig. 1.**
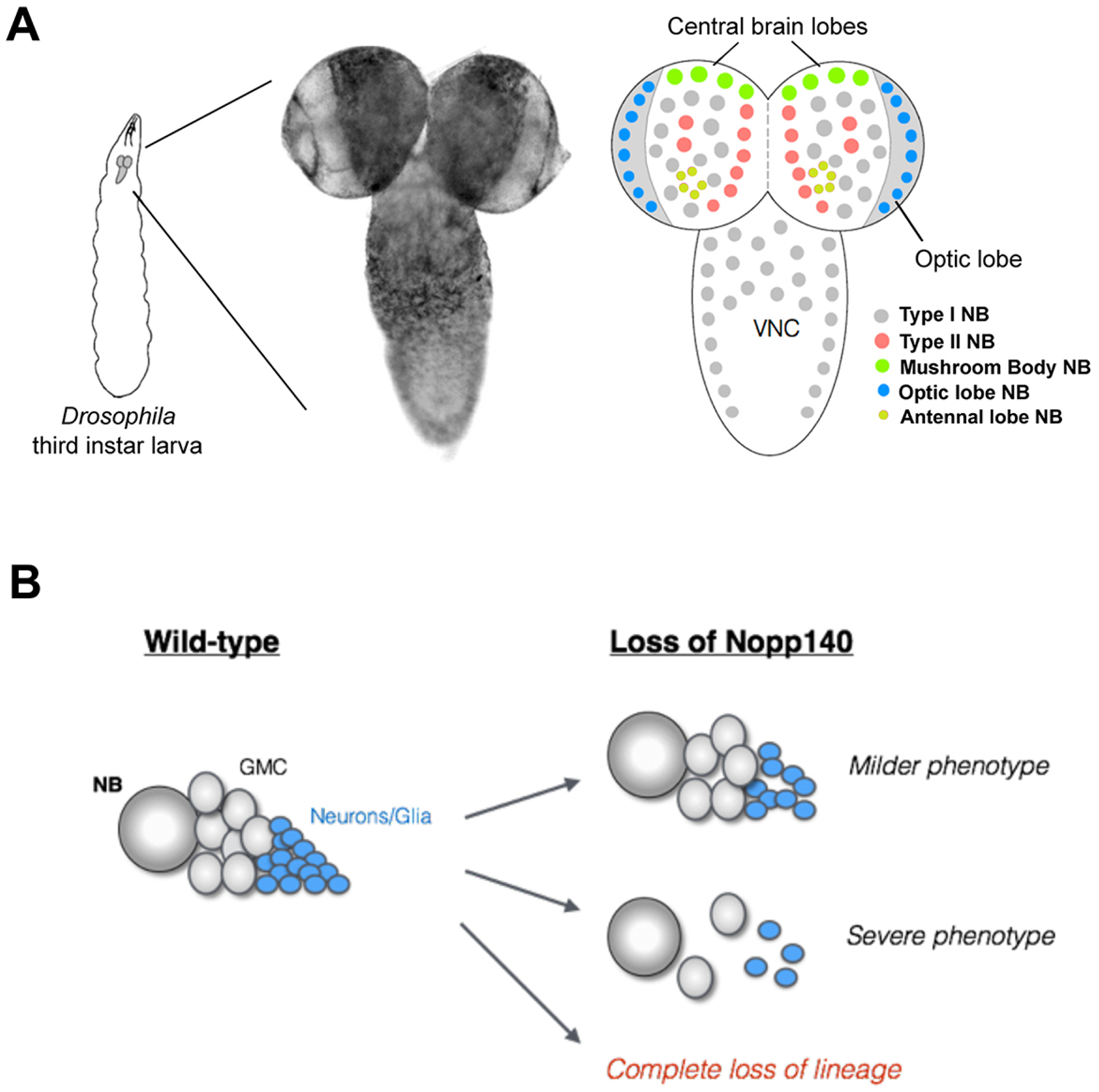
Anatomy of the *Drosophila* larval brain and overall hypothesis. **A** Larval brains have two central brain lobes and a ventral nerve cord (VNC). There are roughly five neuroblast (NB) types within the larval brain: Type I (grey), Type II (red), Mushroom body (green), Antennal lobe (neon), and Optic lobe (blue). These NBs are shown in their putative locations within the larval brain. **B** Our hypothesis is that upon nucleolar stress due to loss of Nopp140, different neuroblast lineages exhibit variable phenotypes ranging from a mild to severe loss of lineage progeny cells, to compete loss of the lineage altogether.

## MATERIALS AND METHODS

### Fly stocks

Fly lines used in this study included: *w^1118^* (used as a wild type control, Bloomington stock #3605), the third chromosome balancer stock *w*; Sb^1^/TM3, P{ActGFP}JMR2, Ser^1^* (referred to as *TM3-GFP*, Bloomington stock #4534), *y^1^ M{nos-Cas9.P}ZH-2A w\** (referred to as *nanos-Cas9*, Bloomington stock #54591 provided by Fillip Port and Simon Bullock, MRC Laboratory of Molecular Biology), *w*; P{GawB}OK107 ey^OK107^/In(4) ci^D^, ci^D^ pan^ciD^ sv^spa-pol^* (referred to as *OK107-GAL4*, Bloomington stock #854), w*; *P{wor.GAL4.A}2; Dr^1^/TM3, P{Ubx-lacZ.w^+^}TM3, Sb^1^* (referred to as *worniu-GAL4*, Bloomington stock #56553), *w^1118^; P{GMR37H04-GAL4}attP2* (referred to as *Scabrous* (*Sca*)*-GAL4*, Bloomington stock #49969), *w^1118^; P{y*[*+t7.7*] *w*[*+mC*]*=GMR38F05-GAL4}attP2* (referred to as *Neurotactin (Nrt)-GAL4*, Bloomington stock #49383), *y^1^ w*; P{w^+mC^=UAS-mCD8::GFP.L}LL5, P{UAS-mCD8::GFP.L}2* (referred to as *UAS-mCD8-GFP*, Bloomington stock #5137), *KO121 Nopp140* gene deletion line (He et al., 2015), and the *UAS-TComC4.2 Nopp140* RNAi line (Cui and DiMario, 2007). Flies were maintained in the laboratory at room temperature (22-24°C) on standard cornmeal-molasses medium. All applicable international, national, and/or institutional guidelines for the care and use of animals were followed.

### Homology Directed Insertion of *DsRed* into *Nopp140*

We used CRISPR-Cas9 and homology directed repair to insert the *DsRed* gene into the second exon of the *Nopp140* gene. The CRISPR optimal target finder tool (http://targetfinder.flycrispr.neuro.brown.edu/) provided 271 gRNA target sites, each 20 nt in length excluding the NGG PAM sequence. Among these, six gRNAs had zero off-targets in coding regions of the *Drosophila* genome. The gRNAs were additionally verified to have no off-targets by the TagScan tool (Genome-wide Tag Scanner; https://ccg.epfl.ch//tagger/tagscan.html), and the Cas-OFFinder tool (Bae et al., 2014). Two gRNA targets, gRNA#52 (5’GGGCTTTGCCGGTTCTTCCT<u>CGG</u> on the minus strand of *Nopp140*; with the PAM sequence underlined) and gRNA #99 (5’CAAGTTGGCTCCTGCTAAGA<u>AGG</u> on the plus strand of *Nopp140*), were chosen and used for CRISPR gene editing. Successful CRISPR-Cas9 cleavage at both gRNA target sites would delete 321 bps from the second exon.

To express these gRNAs, sense and anti-sense oligos that included *BbsI* restriction site overhangs were prepared for both gRNAs by Integrated DNA Technologies (IDT; see Table 1 for gRNA sequences). Mixtures of sense and anti-sense oligos for each gRNA were annealed (heated at 95°C for 5 min, and then cooled to room temperature over 1 hr in 1X ligation buffer). The resulting double-strand DNAs were ligated separately into *pCFD3-dU6:3gRNA* at the *BbsI* site. *pCFD3-dU6:3gRNA* was a gift from Simon Bullock (Addgene plasmid # 49410; http://n2t.net/addgene:49410; RRID:Addgene_49410; (Port et al., 2014; Ren et al., 2013). The resulting plasmids are referred to as *gRNA#55* and *gRNA#99* (Fig. 2A).

**Table 1.**
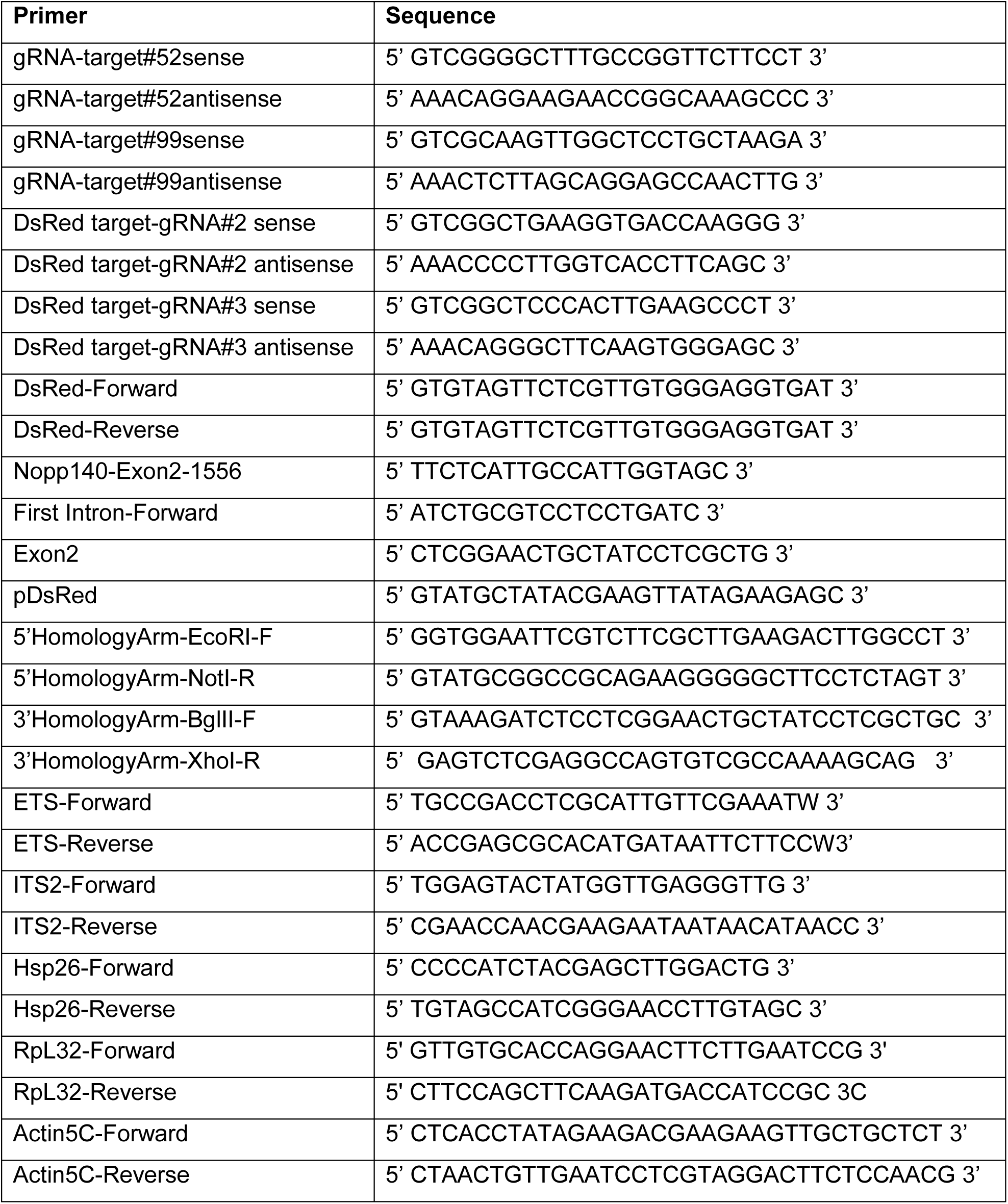
List of primers and their sequences.

**Fig. 2.**
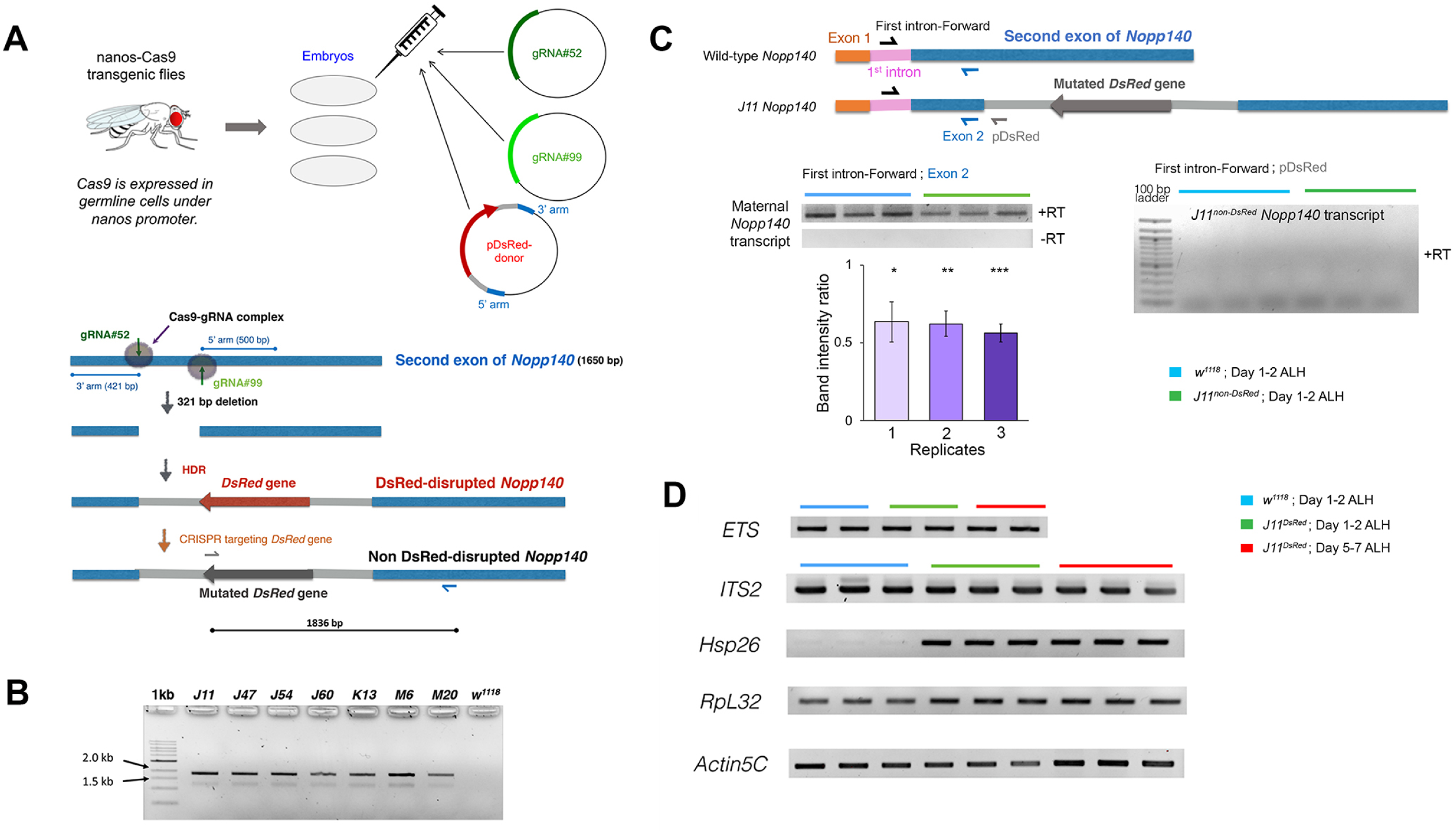
CRISPR-mediated disruption of the *Nopp140* gene and RT-PCR analysis. **A** Three plasmids encoding guide RNAs, gRNA#52 and gRNA#99, or the DsRed protein were injected into embryos from the *nanos-Cas9* fly line. The guide RNAs directed Cas9 cleavage at two specific sites located 321 base pairs apart in the second exon of the *Nopp140* gene (blue bar; 1650 bp total). The *DsRed* gene (red arrow) with flanking plasmid sequences (light grey) and 3’ and 5’ *Nopp140* homology arms inserted into the deletion by homology directed repair (HDR). Seven heterozygous *Nopp140* disruption lines were identified by DsRed expression in adult eyes. The *DsRed* gene was subsequently mutated (dark grey arrow) by CRISPR-mediated mutagenesis. **B** Genomic PCRs using a *DsRed*-specific forward primer (grey) and a downstream *Nopp140*-specific reverse primer (blue half arrow in panel A) verified the *DsRed* insertion within the second exon of *Nopp140* gene in all seven *Nopp140* heterozygous disruption lines (*J11, J47, J54, J60, K13, M6, M20*) with the expected 1836 bp product. The *w^1118^* fly line served as negative control. **C** RT-PCR analyses of *Nopp140* transcript levels in control *w^1118^* and homozygous *J11^non-DsRed^* larvae day 1-2 ALH using the Exon2 reverse primer (blue half arrow) for first strand cDNA synthesis. Subsequent PCRs used the same Exon2 reverse primer and a forward primer specific for the first intron of the *Nopp140* gene. A representative gel is provided. No PCR products appeared in the minus-RT controls. Band intensity ratios (*J11/w^1118^*) were determined for three biological replicates with overall mean +/− SEM of 0.61+/−0.022. Student’s t-test: two-tailed with equal variance for three biological replicates (1, 2, 3) with three PCR technical replicates each, p-values = 0.037*, 0.00081**, 0.0064*** respectively. No RT-PCR product was detected using the pDsRed reverse primer (grey half arrow) in the *J11^non-DsRed^* disruption line. **D** RT-PCR analyses of *ETS, ITS2, Hsp26, RpL32*, and *Actin5C* transcript levels were carried out in control *w^1118^* larvae at day 1-2 ALH and in homozygous *J11^DsRed^* larvae at two time points, day 1-2 and 5-7 ALH using gene-specific reverse primers. Three biological replicates were performed. For each replicate, total RNA was extracted from ~300 *w^1118^* and 150-300 homozygous *J11* larvae. PCR reactions were carried out in triplicate for each first strand cDNA. Representative gels are shown.

To mark the disrupted *Nopp140* gene, the *DsRed* gene was inserted at the Cas9-mediated deletion site by Homology Directed Repair (HDR). We used the donor plasmid, *pDsRed-attP* which was a gift from Melissa Harrison, Kate O’Connor-Giles, and Jill Wildonger (University of Wisconsin-Madison) (Addgene plasmid # 51019; http://n2t.net/addgene:51019; RRID:Addgene_51019; Gratz et al., 2014). We followed general guidelines (Gratz et al., 2014) to insert the homology arms into the multiple cloning sites available on either side of the *DsRed* gene in *pDsRed-attP*. The 5’ and 3’ homology arms from the *Nopp140* second exon were prepared by PCR using forward and reverse primers listed in Table 1. These homology arms flank the 321 bp deletion region described in the preceding paragraph (Fig. 2A). We first inserted the 421 bp 3’ arm into *pDsRed-attP* at the BglII and XhoI sites upstream of the *DsRed* gene, and then inserted the 500 bp 5’ arm at *NotI* and *EcoRI* sites downstream of the *DsRed* gene. The orientation of the homology arms relative to *DsRed* should insert the *DsRed* sequence by HDR such that transcription of *DsRed* is in the opposite direction relative to transcription of the *Nopp140* gene (Fig. 2A). The final plasmid is referred to as *pDsRed-Dono*r.

### NHEJ disruption of *DsRed* gene inserted within *Nopp140* second exon

To mutate the *DsRed* gene within the *Nopp140* gene in the *J11 DsRed* fly line, we used Cas9 endonuclease expressed from the *pBS-Hsp70-Cas9* plasmid, a gift from Melissa Harrison, Kate O’Connor-Giles, and Jill Wildonger (Addgene plasmid # 46294; http://n2t.net/addgene:46294; RRID:Addgene_46294; Gratz et al., 2013). To find gRNA target sites within the *DsRed* gene, we again used the CRISPR optimal target finder tool which yielded 38 gRNA target sites that were 18-nt in length. Twelve of the 38 gRNA targets had no matches to the *Drosophila* genome. Among the twelve gRNA targets, we chose gRNA#2 (5’GCTGAAGGTGACCAAGGG<u>CGG</u> on the plus strand of *DsRed*) and gRNA#3 (5’GCTCCCACTTGAAGCCCT<u>CGG</u> on the minus strand of *DsRed*). Sense and anti-sense oligos for each gRNA target site were prepared by IDT (see Table 1 for sequences). Each double stranded DNA encoding the respective gRNAs was separately ligated into the *pCFD3-dU6:3gRNA* plasmid at the *BbsI* restriction site following the same procedures described above for the preparation of *gRNA#52* and *gRNA#99* plasmids. The resulting plasmids for *DsRed* gene mutagenesis are *gRNA#2* and *gRNA#3*.

### *Drosophila* embryo injections

All plasmids used for embryo injections were extracted from transformed *E. coli* cells using a plasmid Midiprep kit from ThermoFisher Scientific. To disrupt the *Nopp140* gene, the plasmid injection mixture contained 15 ng/μL of *gRNA#52*, 15 ng/μL of *gRNA#99*, and 230 ng/μL of *pDsRed-Donor*. The mixture was injected into homozygous *nanos-Cas9* transgenic embryos. To disrupt the *DsRed* gene, the CRISPR injection mixture contained 75 ng/μL of *gRNA#2*, 75 ng/μL of *gRNA#3*, and 350 ng/μL of *pBS-Hsp70-Cas9*. This mixture was injected into *J11 DsRed/TM3-GFP* embryos. All injections were performed by GenetiVision Corporation (Houston, TX).

### PCR verification of Homology Directed Cas9-mediated donor sequence insertion

Approximately 30 healthy well-fed adults were homogenized in 100 mM Tris-HCl (pH 7.5), 100 mM EDTA, 100 mM NaCl, and 0.5% SDS, followed by 30 min incubation at 70°C. Genomic DNA was precipitated in a 1:2 ratio of 5 M KOAc : 6 M LiCl on ice for 10 min, followed by phenol-chloroform purification and ethanol precipitation. PCR reactions contained 20-70 ng of genomic DNA, 0.40 μM of each primer, 0.20 mM of each dNTP, 0.50 mM of MgCl_2_, 1 X Phusion GC Buffer, and 0.40 unit of Phusion high-fidelity DNA polymerase (M0530S, New England BioLabs). Amplification was performed in a BIO-RAD C1000 Thermal Cycler (cycling conditions: 32 cycles of denaturation for 30 sec at 95°C, annealing for 30 sec at 62°C, and elongation at 72°C for 1 min 20 sec). Primers used for PCR verification were DsRed-Reverse and Nopp140-Exon2-1556. Their sequences are provided in Table 1.

### Sequence analyses

PCR products were extracted from agarose gels using phenol-chloroform, ethanol precipitated, and then sequenced using a BigDye Terminator Cycle Sequencing kit v.3.1 and an ABI 3130XL Genetic Analyzer (Applied Biosystems). Sequencing primers are indicated wherever the sequence reads are provided. Sequences were analyzed and aligned using CLC Sequence Viewer (QIAGEN Bioinformatics).

### RT-PCR analysis

Larvae at day 1-2 after larval hatching (ALH) or day 5-7 ALH were collected from well-yeasted grape juice plates, placed into an Eppendorf tube, and rinsed with distilled water to remove yeast and other debris. Total RNA was extracted from wild-type or *Nopp140−/−* larvae using TRIzol (Invitrogen) according to the manufacturer’s recommendations. First-strand cDNA synthesis was performed using M-MuLV Reverse Transcriptase (NEB M0253S) according to manufacturer’s recommendations with either oligo(dT) primers or gene-specific reverse primers (same as the reverse primers used in PCR). Oligo(dT) primers were used to synthesize the first-strand cDNA of *Hsp26*, *RpL32*, and *Actin5C*. Gene-specific reverse primers were used for the *ETS and ITS2* regions of pre-ribosomal RNA. Specific forward and reverse PCR primers are described in Table 1.

### Immunostaining and fluorescence microscopy

Larval brains and other tissues were dissected directly into fixation Buffer B, pH 7.0-7.2 (16.7 mM KH_2_PO_4_/K_2_HPO_4_, 75 mM KCl, 25 mM NaCl, 3.3 mM MgCl_2_) (de Cuevas and Spradling, 1998) with 2% paraformaldehyde (from a freshly prepared 10% stock). Tissues were fixed for 30-35 min total starting from the point when the dissection commenced. All washings were done with PBS with 0.1% TX-100 detergent. The blocking solution was 3% BSA prepared in PBS with 0.1% TX-100 which was also used for preparing dilutions of primary and secondary antibodies. In all cases, tissues were incubated in the primary antibody overnight at 4°C on a shaker, and in the secondary antibody for 4 hr at 4°C on a shaker. Primary antibodies included the polyclonal guinea pig anti-Nopp140-RGG (Cui and DiMario, 2007) used at 1:100, a rat monoclonal anti-Deadpan (abcam, 195173, stock 1 mg/ml) used at 1:250, the mouse monoclonal anti-fibrillarin mAb 72B9 (Reimer et al., 1987; hybridoma supernatant used without dilution), the mouse monoclonal anti-prospero (deposited at the DSHB by C.Q. Doe; DSHB Hybridoma Product: Prospero MR1A) used at 1:50, and the mouse monoclonal anti-discs large (dlg) (deposited at the DSHB by C. Goodman; DSHB Hybridoma Product: 4F3 anti-discs large) used at 1:30. Secondary antibodies included the Alexa Fluor 546 conjugated goat anti-rat (A-11081, ThermoFisher Scientific) used at 1:1000, the Alexa Fluor 594 conjugated goat anti-guinea pig (A-11073, ThermoFisher Scientific) used at 1:500, and the Dylight 488 conjugated goat anti-mouse (35503, ThermoFisher Scientific) used at 1:500. Tissues were counter-stained with 4’,6-diamino-2-phenylindole (DAPI, Polysciences) at 1 μg/mL. To image the tissues, we used either a conventional fluorescence microscope, a Zeiss Axioskop equipped with a SPOT RTSE digital camera, or a Leica SP8 Confocal Microscope equipped with the White Light Laser system in the Shared Instrumentation Facility (SIF) at Louisiana State University.

### EdU labeling

For 5-ethynyl-2-deoxyuridine (EdU) labeling, larval brains were dissected in PBS (without any detergent or azide), and within 5 min of dissection, the brains were incubated with 20 μM EdU in PBS for 30 min or 2 hr at room temperature. The tissues were then fixed in Buffer B with 2% paraformaldehyde (described above) for 30 min at room temperature. EdU incorporated into S-phase cells was detected by a Click-iT Alexa Fluor 488 EdU imaging kit (Invitrogen) according to the manufacturer’s recommendation. EdU was also detected by Alexa Fluor 594-Azide (Product No.1295, AF 594 Azide from Click Chemistry Tools) used with the reagents provided by Invitrogen Click-iT EdU imaging kit. Following EdU labeling, the larval brains were immunostained with antibodies followed by DAPI counterstaining.

### Determination of nuclear area

The 2D confocal images of the *Nopp140−/−* and wild-type larval brains at day 2-3 ALH were analyzed using Fiji software. After setting scale for each image, the free hand selection tool was used to draw outlines of each nucleus, and the nuclear area was subsequently recorded. Deadpan-stained larval brains were used to determine the nuclear area of neuroblasts. Neuronal nuclear area was obtained from the DAPI-stained larval brains. The nuclear areas were plotted into a box-scatter plot using Microsoft Excel, and a Student’s t-test (one-tailed) was performed on the data.

## RESULTS

### CRISPR for homology directed repair (HDR) to disrupt the *Nopp140* gene

We used CRISPR-Cas9 to delete a target sequence of 321 bps from the second exon of the *Nopp140* gene. A cocktail of two *gRNA* plasmids and the *DsRed-Donor* plasmid was injected into embryos homozygous for the *nanos*::*Cas9* transgene (Fig. 2A). The gRNAs directed the Cas9-mediated deletion, and HDR inserted the *DsRed* gene across the deletion (Fig. 2A). *DsRed* then served as a selectable marker for the disrupted *Nopp140* gene; it was expressed from the *3xP3* eye promoter which is normally active in the entire embryonic and larval brain, Bolwig’s organ, hind gut, anal pads, and adult eyes. We recovered seven independent *Nopp140* disruption lines (*J11, J47, J54, J60, K13, M6, M20*) using the red fluorescence eye phenotype. Each of the seven *Nopp140* disrupted chromosomes was maintained over the *TM3-GFP* balancer chromosome which carries a wild type copy of *Nopp140*. The *DsRed* insertion was verified by genomic PCRs (Fig. 2B). The expected 1836 bp PCR product was amplified in all seven *Nopp140* insertion alleles, with *w^1118^* acting as a wild type negative control (Fig. 2A,B). Among the seven lines initially recovered, *J11^DsRed^/TM3-GFP* was backcrossed with the *Sb^1^*/*TM3-GFP* fly line for at least six generations to eliminate possible off-target mutations in the *J11^DsRed^* line.

The *GFP* reporter gene on the *TM3* balancer chromosome is expressed in a small cluster of larval midgut cells that are easily identifiable. Therefore, with *inter se* crosses of *J11^DsRed^/TM3-GFP* stock flies, we hand-selected larvae that were homozygous for *J11^DsRed^*, but selected against sibling larvae heterozygous for *J11^DsRed^/TM3-GFP* with prominent GFP signals in their midgut.

To conduct multi-channel immunofluorescence of the *Drosophila* brain, we again used CRISPR-Cas9 but now with Non-Homologous End Joining (NHEJ) to disrupt the *DsRed* gene inserted in the *J11^DsRed^* allele. A cocktail of two gRNA plasmids and the *pBS-Hsp70-Cas9* vector injected into *J11^DsRed^/TM3* embryos produced several independent fly lines with mutations in *DsRed* (Supplementary Fig. S1). We sequenced the second exon region in two of these lines, *A5* and *A7*, and verified that each had a short deletion at the gRNA#3 target site within the *DsRed* gene (Supplementary Fig. S1). The *A7-J11^non-DsRed^* fly line was again back-crossed six times to deplete any possible off-site targets. Either the original *J11^DsRed^* fly line or its derived A7*-J11^non-DsRed^* line was used for the experiments described below.

We next performed RT-PCR analyses to test if the disrupted *Nopp140* gene was transcribed in homozygous *A7-J11^non-DsRed^* larvae (day 1-2 ALH) (Fig. 2C). The reverse primer referred to as *pDsRed* in Fig. 2C annealed to the *pDsRed-attP* plasmid sequence a few base pairs downstream of the junction between the *Nopp140* second exon and the *DsRed* donor sequence. No transcripts containing *DsRed-att* sequences were detected in the total RNA samples prepared from homozygous *A7-J11^non-DsRed^* larvae 1-2 day, similar to the *w^1118^* sample that served as a negative control (Fig. 2C, but see below for a positive control). Nonsense-mediated decay (NMD) would likely degrade *Nopp140* pre-mRNAs transcribed from the disrupted gene as they would likely contain premature stop codons within the *pDsRed-att* sequences, or these pre-mRNAs may be improperly/incompletely spliced (Garneau et al., 2007). This lack of RT-PCR products eliminates the likelihood of a dominant-negative effect due to the production of truncated Nopp140 proteins encoded by the disrupted *Nopp140* gene. In summary, hand-selected larvae homozygous for the *Nopp140 J11^non-DsRed^* allele provide a null genotype (*Nopp140−/−*) systemic throughout the larvae.

We next determined if there were maternal wild type *Nopp140* transcripts present in the RNA preparations isolated from the same homozygous *A7-J11^non-DsRed^* 1-2 day ALH larvae used for the RT-PCRs described in the preceding section. We performed these second RT reactions using a reverse primer (*Exon2*, blue in Fig. 2C) that anneals to the *Nopp140* second exon a few base pairs upstream of the junction between the *Nopp140* second exon and the *DsRed* donor sequence. Since *Nopp140* transcripts harboring *DsRed* sequences were undetectable in these larvae, first strand cDNAs primed with *Exon2* should indicate the presence of maternal *Nopp140* transcripts in homozygous *J11^DsRed^* larvae, and thus serve as a positive control for the initial RT-PCRs that showed an absence of *DsRed-att*-containing transcripts. These second RT-PCRs showed that maternal *Nopp140* transcripts were indeed present in the *Nopp140−/−* larvae at day 1-2 ALH. The abundance of maternal *Nopp140* transcripts in the *Nopp140−/−* larvae was about half that seen in wild type larvae, suggesting that both maternal and zygotic *Nopp140* transcript pools exist in early wild type larvae (Fig. 2C).

As additional controls (Fig. 2D), RT-PCR analyses of the External Transcribed Spacer (*ETS*) and the Internal Transcribed Spacer 2 (*ITS2*) sequences within pre-rRNA showed that their levels were unaffected in homozygous *J11^DsRed^* larvae at day 1-2 ALH and at day 5-7 ALH. This indicates that loss of Nopp140 had no effect on rDNA transcription, which agreed with our earlier observations with a *pBac*-generated *Nopp140^KO121^* deletion line (He et al., 2015). Furthermore, *Hsp26* transcript levels were upregulated in homozygous *J11^DsRed^* larvae at both day 1-2 and day 5-7 ALH, whereas the wild-type larvae had almost undetectable levels of *Hsp26* transcript (Fig. 2D). Overexpression of *Hsp26* in homozygous *J11^DsRed^* larvae as early as day 1 ALH indicated a cellular stress response due to the effects of Nopp140 loss (e.g., Wang et al., 2004). As final controls, we accessed *RpL32* and *Actin5C* transcript levels: while *RpL32* transcript levels remained unchanged between the wild-type and homozygous *J11^DsRed^* samples and between biological replicates, *Actin5C* transcript levels fluctuated slightly within the samples and between biological replicates for reasons that remain uncertain.

### Maternal Nopp140 protein is reduced in early *J11 Nopp140−/− larval* brains

Since the RT-PCR analyses showed that maternal *Nopp140* transcripts persisted in the homozygous *J11^DsRed^* larvae at day 1-2 ALH, we wanted to test if the Nopp140 protein could be detected in their brain and gut tissues as well. To do this, we immunostained homozygous *A7-J11^non-DsRed^* larvae and wild-type larvae with an antibody directed against Nopp140-RGG, one of the two Nopp140 isoforms in *Drosophila*. This antibody was raised against a synthetic peptide, the sequence of which is unique to the carboxyl tail region of Nopp140-RGG (see Cui and DiMario, 2007). An antibody directed against the carboxyl terminus of the other isoform, Nopp140-True, has proven much weaker, and was not used here. At day 1-2 ALH, the anti-Nopp140-RGG antibody labeled nucleoli in homozygous *J11 ^non-DsRed^* larval brain and midgut, but at lower levels compared to the same wild type tissues (Fig. 3, panels a-d for brain, e-h for gut tissue). The homozygous *A7-J11^non-DsRed^* larval brains had fewer and smaller-sized nucleoli compared to nucleoli in wild-type brains at day 1-2 ALH (Fig. 3, compare panels a and c). Four large-sized nucleoli per brain lobe were routinely detected in the anterior of wild-type larval brains, and we speculated these were the Mushroom Body (MB) neuroblasts that do not undergo quiescence, but continue to divide throughout the embryo-to-larva transition (arrow in Fig. 3a). However, we did not observe this preferential labeling in homozygous *A7-J11 ^non-DsRed^* brains. By day 4-5 ALH, nucleolar labeling by anti-Nopp140-RGG was noticeably reduced in homozygous *A7-J11 ^non-DsRed^* larval brains and gut tissues as compared to the wild type tissues (Fig. 3, panels i-l for brain, m-p for midgut). These results indicated that at least the Nopp140-RGG isoform encoded presumably by maternal transcripts persisted in the first two days of homozygous *A7-J11 ^non-DsRed^* larval development, but then diminished in most cells as these larvae aged.

**Fig. 3.**
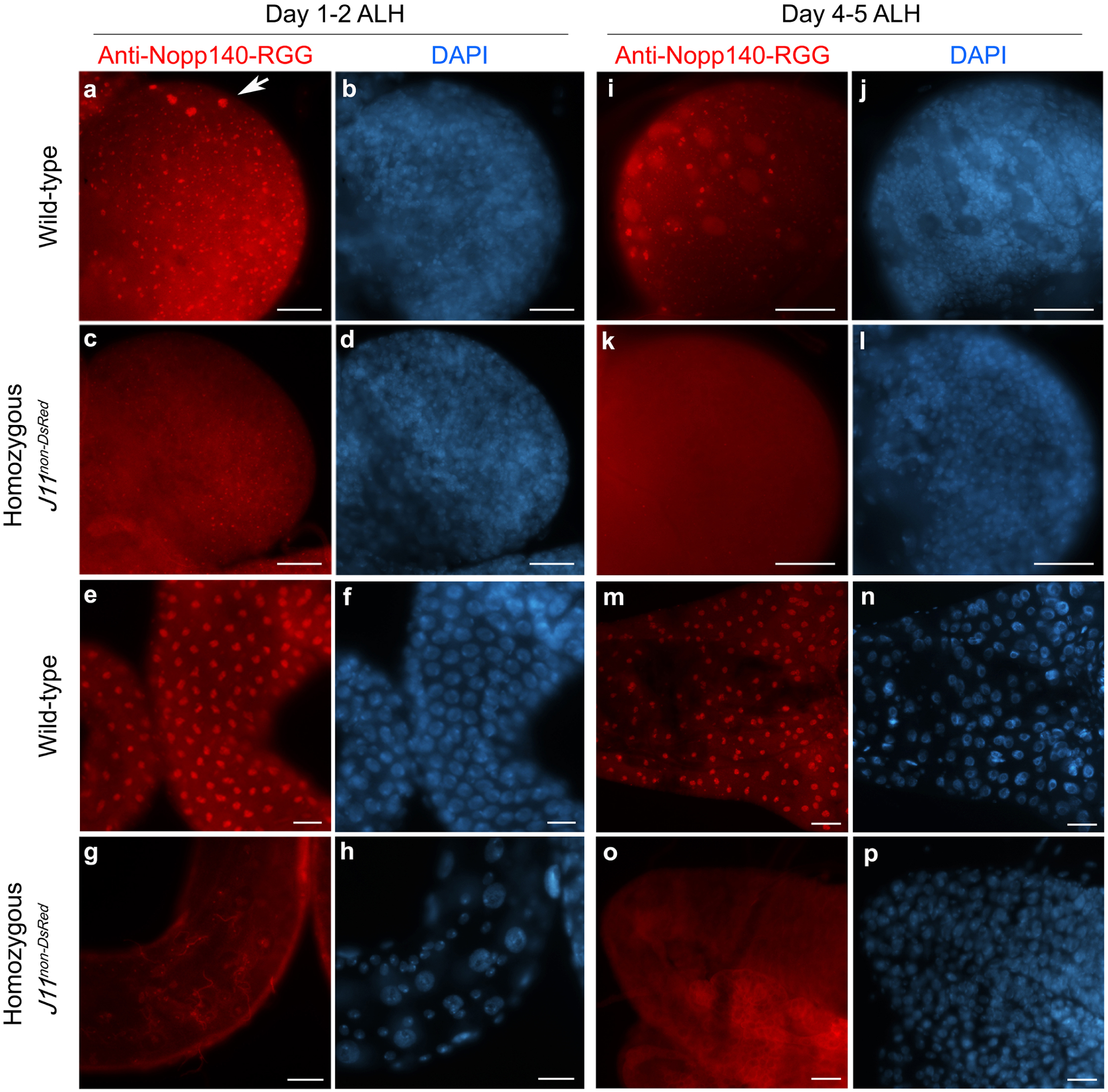
Maternal Nopp140 protein is reduced in early *J11 Nopp140−/− larval* brains. Central brain lobes and mid-gut tissues from *w^1118^* control (a-j, e-n) and homozygous *J11^non-DsRed^* larvae (c-l, g-p) at day 1-2 (a-h) and day 4-5 (i-p) ALH were immunostained with anti-Nopp140-RGG. Arrow in **panel a** indicates four neuroblasts per wild type brain lobe with large nucleoli labeled with anti-Nopp140-RGG. n=19 (*w^1118^*); n=23 (homozygous *J11^non-DsRed^*); 2 technical replicates. Scale bars: 10 μm in panels a-f, m-p; 25 μm in panels g-l

### Embryonic and larval survivability with complete or partial elimination of Nopp140

The *Nopp140* disruption lines were maintained using the third chromosome balancer, *TM3*, which carries a wild type *Nopp140* gene. Embryos homozygous for *TM3* are non-viable, hence *inter se* crosses within the *J11 ^DsRed^/TM3* fly stock should produce 50% *Nopp140^DsRed^/TM3* larvae and 25% homozygous *J11 ^DsRed^* larvae (the number of hatched larvae ÷ total number of eggs collected). However, if the disrupted *Nopp140* gene causes embryonic lethality, we would expect frequencies less than 50% and 25%, respectively. We found that only 20.8% of total eggs developed into larvae that were *J11 ^DsRed^/TM3* versus the expected 50% (Fig. 4A), and only 7.1% of the total eggs developed into larvae that were homozygous for *J11^DsRed^* versus the expected 25% (Fig. 4A). These data indicated that loss of Nopp140 leads to partial embryonic lethality not only for the *homozygous J11^DsRed^* genotype, but more interestingly for the heterozygous *J11^DsRed^/TM3* genotype. The observation indicated for the first time that the *Nopp140-/+* genotype exhibits haplo-insufficiency in *Drosophila*, similar to the *Tcof1-*/+ genotype in the human Treacher Collins Syndrome.

**Fig. 4.**
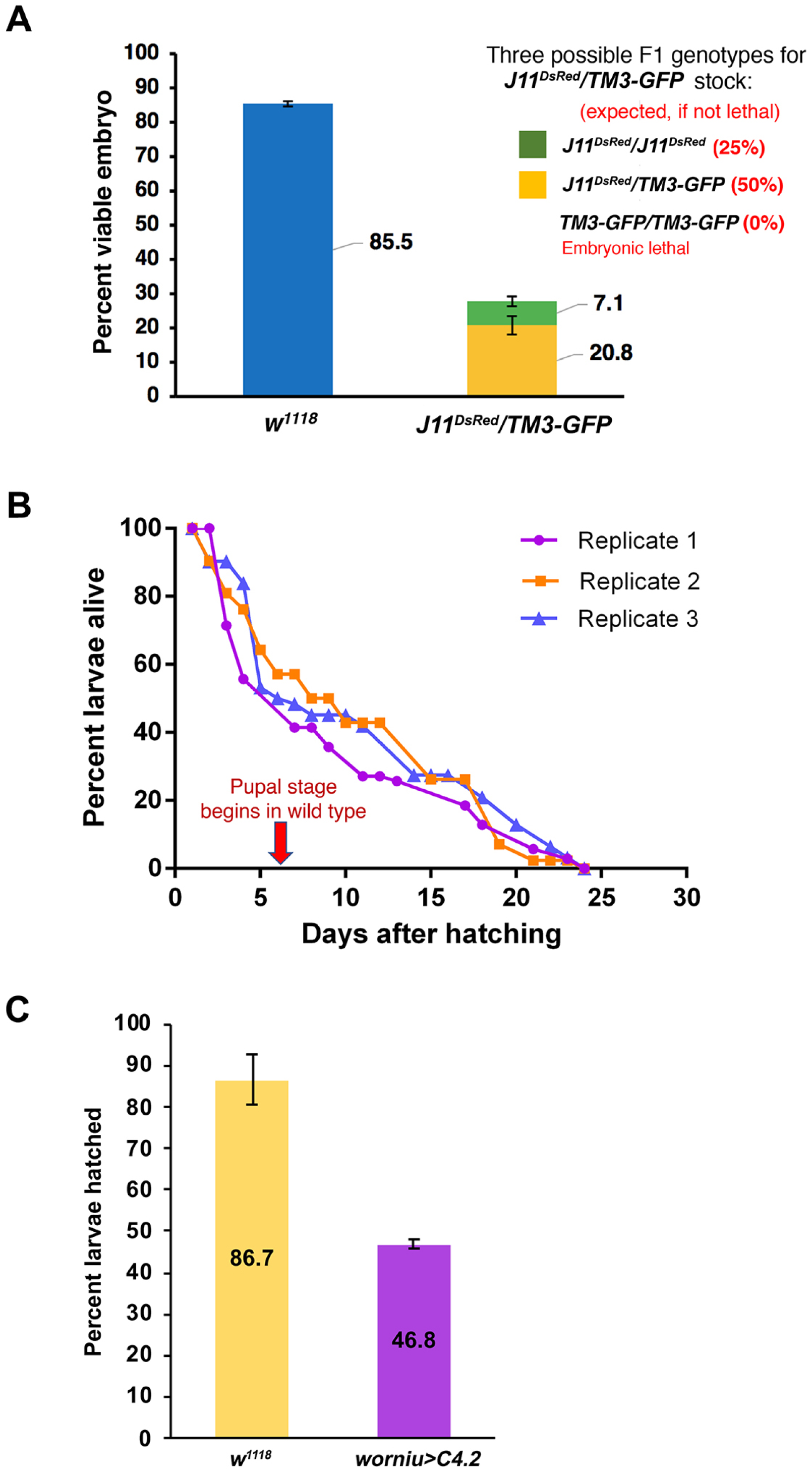
Embryonic and larval survival upon complete or partial loss of Nopp140. **A** Survival assays were performed for homozygous *J11^DsRed^* or heterozygous *J11^DsRed^/TM3-GFP* embryos, and for *w^1118^*control embryos. Freshly laid eggs were collected from well-yeasted juice plates (n = 2; total number of embryos per replicate for *w^1118^*: 200 and 299, and for *J11^DsRed^/TM3-GFP* stock: 230 and 111). The number of hatched larvae were logged for the next two days, and the percent viable embryos was determined. Data shows the number of larvae hatched divided by the total number of embryos collected X 100%. **B** Plot shows three replicates (number of larvae per replicate: 70, 42, 62) of survival assay for homozygous *J11^DsRed^* larvae. Newly hatched larvae were collected from a well-yeasted juice plates, and the number of living larvae were recorded in the following days until all larvae had perished. **C** Embryonic lethality and larval survivability upon Nopp140 depletion by RNAi expression using the *worniu::GAL4* driver (specific for all embryonic and larval neuroblasts) and *UAS::TComC4.2* (Nopp140-RNAi line; Cui and DiMario 2007). Compared to 86.7% of the *w^1118^* embryos, only 46.8% of the collected embryos with Nopp140 depletion (*worniu-GAL4>C4.2*) hatched and developed into 3rd instar larvae, after which all larvae developed into adults (not shown). n = 3; total number of embryos collected per replicate for each genotype: 200, 265, and 330; Student’s t-test: two-tailed with unequal variance, p-value = 0.0069

We earlier described growth arrest and lethality in second instar larvae that were homozygous for our original *pBac*-mediated *Nopp140^KO121^* deletion (He et al., 2015). Because of the particular *pBac* elements available at the time, we had to delete the 3’ end of the downstream gene, *P5CDh1* (He and DiMario, 2011), and this constantly forced us to control for the carboxyl truncation in the protein product when assessing the loss of Nopp140. Here, we assessed survivability of larvae homozygous for *J11^DsRed^*. Similar to our earlier findings (He et al. 2015), we found that ~50% of the homozygous *J11^DsRed^* larvae died by day 6 (which is when the pupal stage normally begins) (Fig. 4B). The remaining 50% remained as second instar larvae; they failed to grow or molt. The number of surviving homozygous *J11^DsRed^* larvae dwindled over time, but interestingly, some lingered up to day 24 (Fig. 4B).

We also depleted Nopp140 using the UAS-GAL4 system to express siRNAs. In the past we showed that *daughterless::GAL4*>*UAS::TComC4.2* depleted ~70% of the *Nopp140* transcripts (Cui and DiMario, 2007). Using the neuroblast-specific *worniu::GAL4* driver (*worniu-GAL4*>*UAS::TComC4.2*), we found embryonic survivability was ~46%, while the wild type embryo survival rate was ~86% (Fig. 4C). Interestingly, the surviving *worniu::GAL4*>*UAS::TComC4.2* larvae developed into viable and fertile adults. While the *worniu* promoter is active in all embryonic and larval neuroblasts, its peak expression is in 6-12 hr embryos, perhaps explaining the survivability of nearly half the *worniu::GAL4*>*UAS::TComC4* embryos beyond this embryonic stage.

### Brain hypoplasia upon nucleolar stress

We found that larval brain development was severely impaired upon loss of Nopp140 either by gene disruption (i.e., homozygous *J11 ^DsRed^)* or by neuron-specific RNAi depletion. During the early larval stage (day 1-2 ALH), homozygous *J11^DsRed^* brains were morphologically comparable in size to brains from newly hatched wild-type larvae. The mutant’s brain continued to grow from day 3-6 ALH, but more slowly compared to wild-type larval brains (Fig. 5A). Beyond day 5-6 ALH, homozygous *J11^DsRed^* larval brains failed to grow. This was similar to what we saw in our original *Nopp140^KO121^* deletion (He et al., 2015) (Fig. 5A). Likewise, brain growth was impaired in larvae upon RNAi-mediated depletion of Nopp140 using a pan-neuronal *GAL4* driver *(Neurotactin::GAL4>UAS::TComC4.2*) (Fig. 5B).

**Fig. 5.**
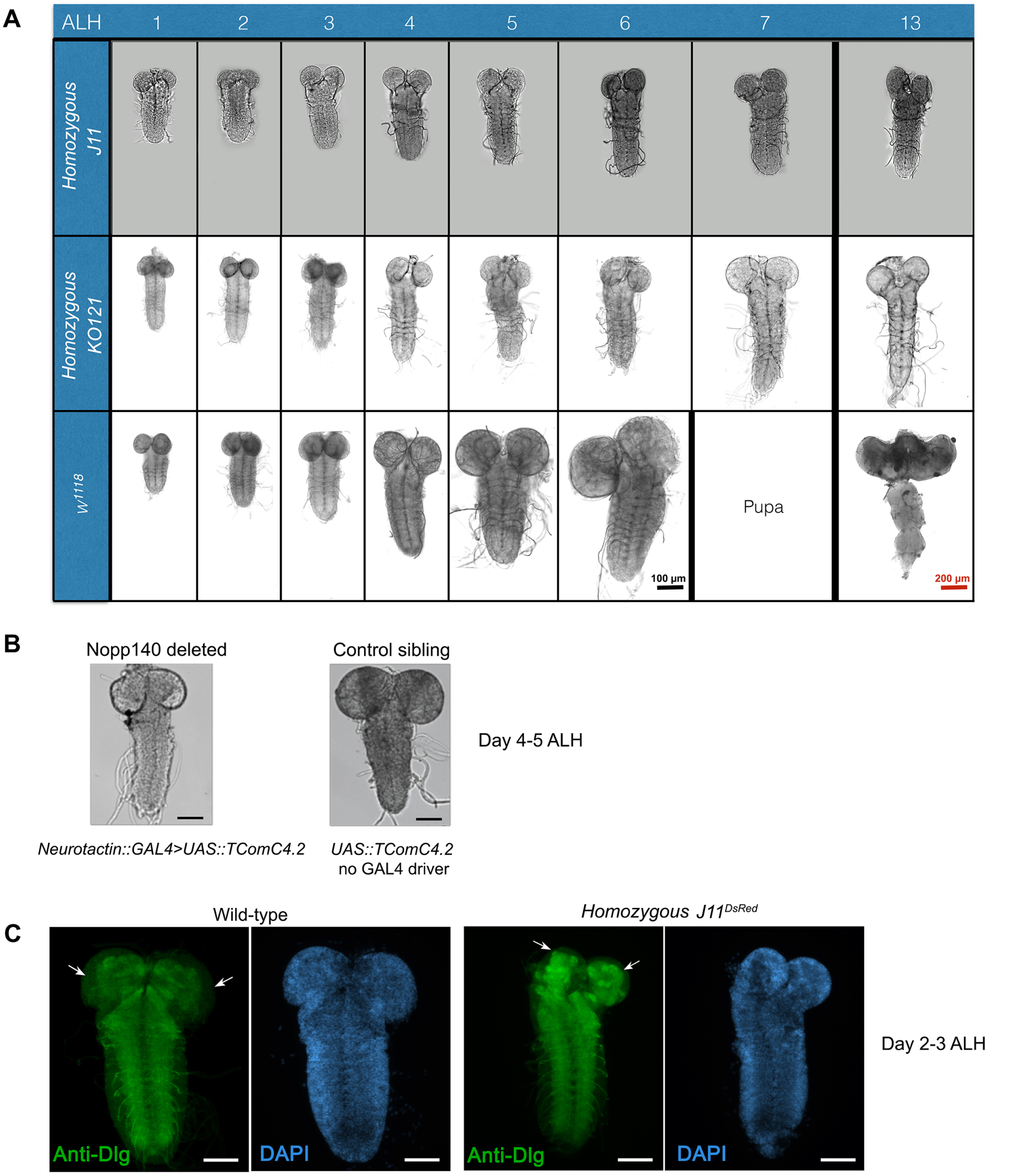
*Drosophila* larval brain development is impaired under nucleolar stress induced by the loss of Nopp140. **A** Larval brain development in homozygous *KO121* (*Nopp140* gene deletion, He et al. 2015), homozygous *J11* (CRISPR-mediated *Nopp140* disruption), and wild-type (*w^1118^*) starting at day 1 after larval hatching (ALH) until day 7 and day 13 ALH. Homozygous *KO121* and *J11* larval brains at day 13 ALH are shown, but wild-type individuals have developed into adults by day 13 ALH, hence an adult fly brain is shown. **B** RNAi-depletion of Nopp140 using pan-neuronal GAL4 driver, *Neurotactin (Nrt)::GAL4*, and the *UAS::TComC4.2* (Nopp140 RNAi line) resulted in impaired larval brain development similar to that seen in Nopp140 homozygous deletion background. Representative larval brains from three independent crosses at day 4-5 ALH comparing Nopp140**-**depleted brains with control sibling brains (not expressing RNAi) are shown. Scale bar: 100 μm **C** Conventional fluorescence images of the neuropil immunostained with antibody against Discs large (Dlg; green) in second **i**nstar *w^1118^* control larvae and homozygous *J11^DsRed^* larvae at day 2-3 ALH. White arrows show unstained peripheral cell body layers which are reduced in homozygous *J11^non-DsRed^* larvae. n=15 (wild-type); n=18 (homozygous *J11^non-DsRed^*); >3 technical replicates. Scale bar: 50 μm

To see where growth was interrupted, we immunostained brains from homozygous *J11^DsRed^ larvae* and wild type larvae at day 2-3 ALH with an antibody against *discs large* (anti-Dlg). This antibody stains axon bundles (the neuropil), but not the cell body mass which we counter-stained with DAPI. Neuropils within the two central brain lodes were reduced in homozygous *J11^DsRed^* brains as compared to wild type brains, but there were no observable physical defects in the ventral nerve cord (VNC) neuropil of homozygous *J11^DsRed^* larvae when compared to wild type larvae. Besides the reduced central brain lobe neuropils, we found the cell body mass of the central brain lobe was also reduced in homozygous *J11^DsRed^* brains when compared to wild type brains (Fig. 5C).

### Reduced neuroblast numbers and proliferation upon nucleolar stress

We hypothesized that the hypoplasia in *Nopp140−/−* larval brains was due to either a reduction in NB numbers, a reduction in their proliferative capacity, or both. To assess these possibilities, we first performed a Click-iT EdU labeling assay on living brains. EdU is a thymidine analog which is incorporated into genomic DNA during S-phase of the cell cycle, and hence these cells are committed to cell division. The assay used a 2 hr EdU pulse in wild type and homozygous *A7-J11^non-DsRed^* larval brains at 1, 2-3, and 6 days ALH. After pulse-labeling, brains were fixed with paraformaldehyde, and the EdU residues were fluorescently labeled by Click-iT chemistry. We then immunostained the same brains with anti-Deadpan to visualize the number and distribution of neuroblasts. Deadpan (Dpn) is a neuroblast-specific transcription factor necessary for self-renewal properties.

Anti-Dpn labeling showed that NBs were present in homozygous *J11^non-DsRed^* larval brains from all age groups, however their numbers were consistently reduced compared to the wild-type brains of the same age (Fig. 6; compare homozygous *J11^non-DsRed^* panels a, g, and m with wild-type panels d, j, and p). This suggested that fewer neuroblasts in the homozygous *J11^non-DsRed^* larval brains likely contributed to the observed hypoplasia. Strikingly, in homozygous *J11^non-DsRed^* larvae at day 1 and day 2-3 ALH, we consistently noticed four NBs in each central brain lobe that showed prominent anti-Dpn labeling compared to the surrounding Dpn-stained NBs (Fig. 6; arrows in panels a and g). These four NBs were visible in the wild-type brains as well, but only in brains from day 1 ALH larvae (Fig. 6; arrows in panel d).

**Fig. 6.**
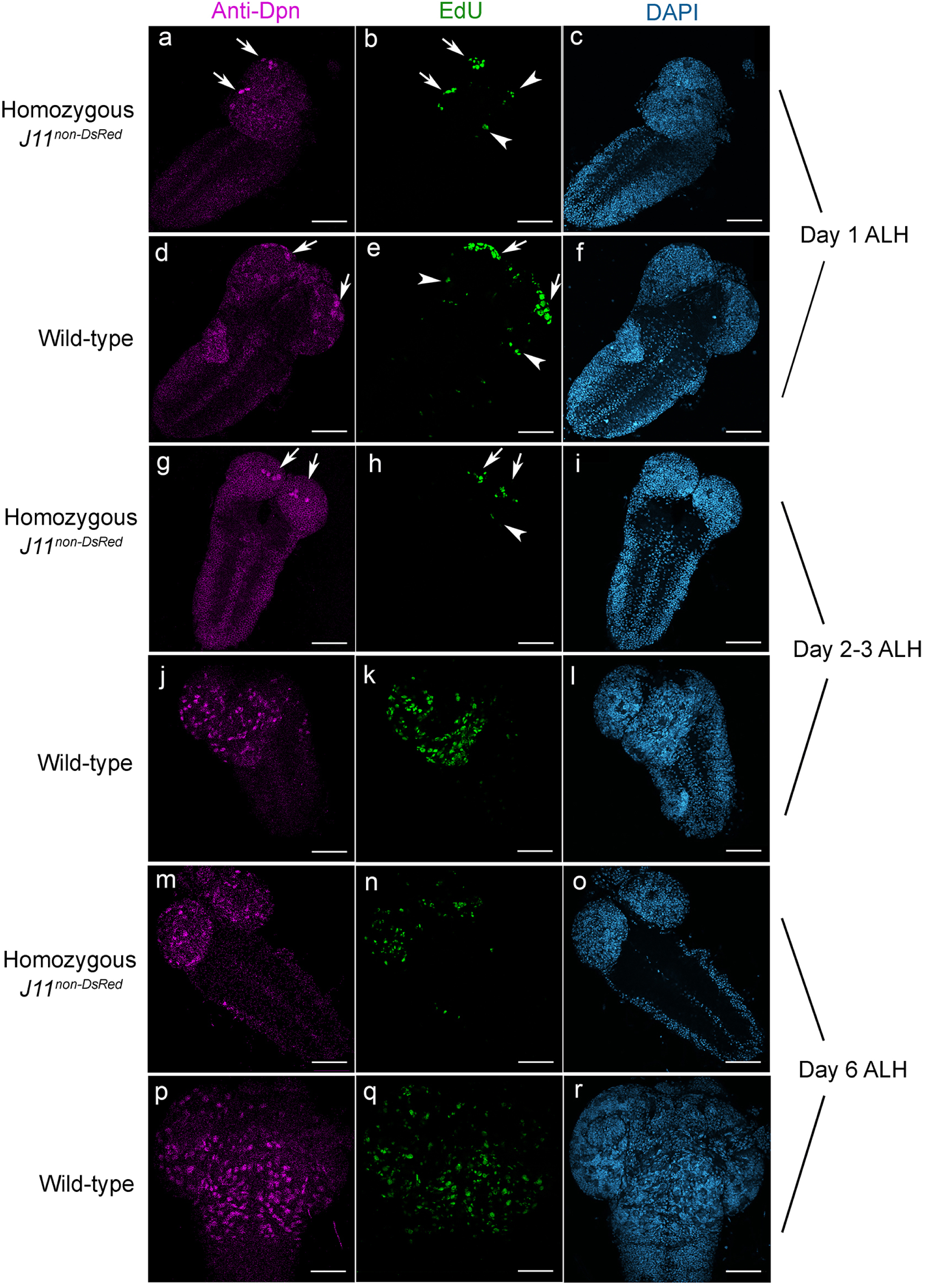
Neuroblast proliferation is reduced upon nucleolar stress. Confocal images of homozygous *J11^DsRed^* and control *w^1118^* larval brains at day 1 (a-f), 2-3 (g-l), and 6 (m-r) ALH are shown after EdU-labeling (Click-iT Alexa Fluor 488) followed by anti-Deadpan (anti-Dpn) immunostaining. Dpn-stained cells (magenta) are neuroblasts. After a 2 hr pulse, EdU-labeled S-phase cells (green) were committed to cell division. Arrows indicate four likely Mushroom Body NBs that were EdU- and Dpn-positive and clustered near the anterior of the central brain (a, b, d, e, g, h). Arrowheads indicate a few EdU-positive cells, likely arising from AL MBs, at the lateral side of the central brain (b, e, h). n=10, 15, and 10 for day 1, 2-3, and 6 respectively for both wild-type and homozygous *J11^DsRed^* sample; 3 technical replicates. Scale bar: 50 μm

EdU labeling displayed NBs and their progeny GMCs that were in S-phase. Overall, homozygous *J11^non-DsRed^* larval brains had fewer EdU-positive cells as compared to the wild-type brains in all three examined age groups (Fig. 6; compare homozygous *J11^non-DsRed^* panels b, h, and n with wild-type panels e, k, and q). The same subset of the EdU-positive cells in both homozygous *J11^non-DsRed^* and wild-type brains were identified as NBs by the anti-Dpn nuclear staining. Other than the four NBs that co-labeled with EdU and anti-Dpn in both homozygous *J11^non-DsRed^* and wild-type larval brains at day 1 ALH, there were also fewer EdU-positive GMCs in the homozygous *J11^non-DsRed^* brains compared to the wild-type brains (Fig. 6; compare *Nopp140−/−* panels a, b with wild type panels d, e). This suggested a slower rate of NB proliferation in the homozygous *J11^non-DsRed^* larval brains compared to the wild-type NBs.

At day 2-3 ALH, we consistently observed only the four anterior NBs that co-labeled with both EdU and anti-Dpn in homozygous *J11^non-DsRed^* brains (Fig. 6, arrows in panels g and h). We predicted that these NBs were the MB NBs based on their location and consistency in number. Wild-type larval brains, however, had more EdU-positive and Dpn-positive cells, suggesting that the majority of NBs had exited quiescence and started to proliferate as expected (Fig. 6, panels j and k). At day 1 ALH, we also noticed other EdU-positive cells in the lateral regions of the central brain lobes from both homozygous *J11^non-DsRed^* and wild-type larvae (Fig. 6, arrowheads in panels b and e). These should be the Antennal Lobe (AL) NBs. They were occasionally detected in central brain lobes of day 2-3 ALH homozygous *J11^non-DsRed^* (Fig. 6, arrowheads in panel h).

These observations indicate that upon nucleolar stress, only a subset of neuroblasts and GMCs proliferate in homozygous *J11^non-DsRed^* brains although at a slower rate, and give rise to lineages that are comparatively smaller than those in wild type brains under non-stressed conditions. Indeed, using an antibody against Prospero, a nuclear marker specific for GMCs and their descendent glia and neurons, we found significantly fewer GMC populations in the homozygous *J11^non-DsRed^* brains than in wild-type brains at day 1-2 and 6-7 ALH (Supplementary Fig. S2). Additionally, we found that the nuclear volumes in the NBs and neurons were noticeably reduced in homozygous *J11^non-DsRed^* larval brains compared to wild-type larval brains at day 2-3 ALH (Supplemental Fig. S3). Thus, upon nucleolar stress, larval brain hypoplasia resulted from the loss of mostly Type I and II neuroblasts, the reduced size of remaining neuroblasts and neurons, and the inability of these neuroblasts to proliferate.

### Mushroom body neuroblasts are resilient to nucleolar stress

To test if the four anterior EdU-positive NBs were in fact MB NBs, we used a MB lineage-specific *GAL4* driver to express a GFP-tagged plasma membrane reporter protein, mCD8-GFP (*OK107::GAL4*>*mCD8::GFP*), and again performed a 30 min co-EdU-labeling in brains from both homozygous *J11^non-DsRed^* and control larvae at day 3 ALH (see the genetic cross scheme in Supplemental Fig. S4). EdU labeling showed many S-phase cells in the wild type larval brains; a subset of these cells located within the mCD8-GFP-positive MB-lineage cell cluster (Fig. 7; panel c). In the homozygous *J11^non-DsRed^* larval brains at day 3 ALH, EdU-positive cells in the anterior region of the CBs were always located within the MB lineage-cell cluster as identified by mCD8-GFP (Fig. 7; panel g). This suggested that the four Dpn-positive and EdU-positive NBs that we observed in homozygous *J11^non-DsRed^* larval brains at day 2-3 ALH (Fig. 6; panel g and h) were indeed MB NBs. The combined results of Figs. 6 and 7 suggest that the MB NBs are more resilient to nucleolar stress induced by the loss of Nopp140 as compared to other NBs within these brains.

**Fig. 7.**
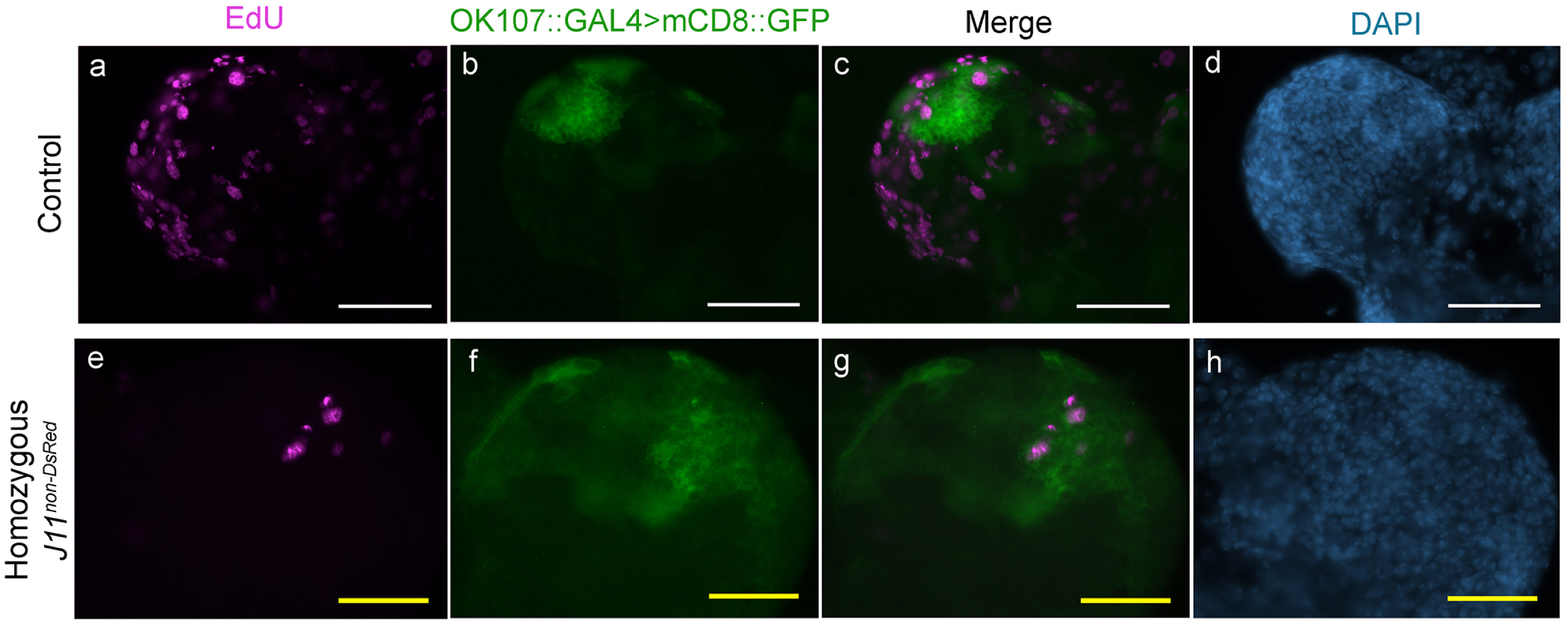
Mushroom Body NBs are resilient to nucleolar stress. Larval brains from control *w^1118^* and homozygous *J11^non-DsRed^* larvae (see the cross scheme in Supplementary Fig S4) at day 3 ALH were used for 30 min EdU pulse labeling (Click-iT Alexa Fluor 594). Merged confocal images show EdU-labeled cells (magenta) nestled within the GFP-labeled MB lineage (green) near the anterior of the central brains. n=12 (control); n=20 (homozygous *J11^non-DsRed^*); 3 technical replicates. Scale bars: 50 μm in panels a-d, 25 μm in panels e-h

### Mushroom body neuroblasts in the *Nopp140−/−* larval brain retain nucleolar fibrillarin

Nopp140 is a chaperone for C/D-box snoRNPs that catalyze 2′-*O*-methylation of pre-rRNA during ribosome biogenesis. Previous work in our lab showed that the C/D-box snoRNP methyl-transferase, fibrillarin, redistributes to the nucleoplasm upon complete loss of Nopp140 in larval tissues homozygous for the *Nopp140^KO121^* gene deletion. Loss of fibrillarin from the nucleoli caused a reduction in 2′-*O*-methylation of pre-rRNA clearly indicating nucleolar dysfunction, even though gross nucleolar morphology and rDNA transcription remained normal (He at al., 2015). Since MB NBs, but not others, continue to divide in *Nopp140−/−* larval brains at day 2-3 ALH, we predicted that MB NBs might retain fibrillarin within their nucleoli, while other NBs and their lineages redistribute fibrillarin to the nucleoplasm. To test this, we immunostained brains from homozygous *J11^non-DsRed^*; *OK107::GAL4>mCD8::GFP* larvae and wild-type *OK107::GAL4>mCD8::GFP* larvae (see the genetic cross scheme in Supplemental Fig. S4) with anti-fibrillarin at day 3 ALH. While anti-fibrillarin stained nucleoli with minimal nucleoplasmic labeling in the wild-type larval brains (Fig. 8), it labeled the nucleoplasm in the majority of brain cells in homozygous *J11^non-DsRed^* larval brains, except for a small number of cells located within the MB-lineage as marked by mCD8-GFP labeling; these cells showed clear nucleolar labeling with anti-fibrillarin, even though there was some nucleoplasmic labeling (Fig. 8). This result indicates that the MB-lineage cells, and not others, are able to retain at least some nucleolar fibrillarin, indicating that their nucleoli are partially functional.

**Fig. 8.**
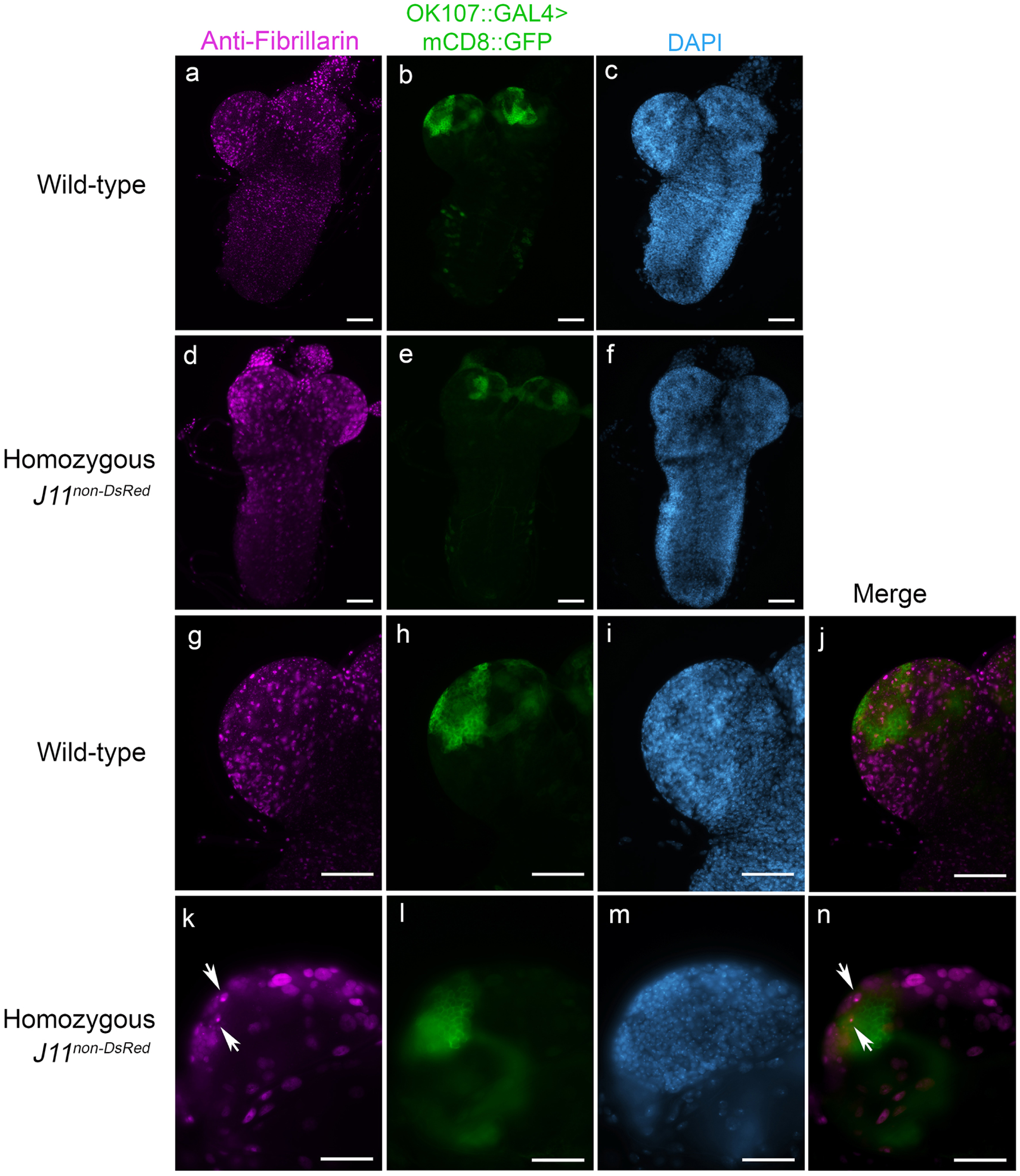
Mushroom Body lineage cells retain nucleolar fibrillarin under nucleolar stress. Larval brains from control *w^1118^* (whole brain in a-c; a central brain lobe in g-j) and homozygous *J11^non-DsRed^* larvae (whole brain in d-f; a central brain lobe in k-n) at day 3 ALH were immunostained with anti-fibrillarin (magenta). Merged images in panels j and n were obtained from confocal images g and h, and k and l respectively. Arrows in the homozygous *J11^non-DsRed^* larval brains (k, n) indicate nucleolar fibrillarin retained in the MB lineage cells marked by cell surface protein, mCD8::GFP (green), whereas fibrillarin is redistributed into the nucleoplasm in surrounding cells. n=10 (wild-type); n=10 (homozygous *J11^non-DsRed^*); 2 technical replicates. Scale bars: 50 μm in panels a-j, 25 μm in panels k-n

### A transcriptomics perspective

With MB NBs apparently retaining fibrillarin, we asked how much Nopp140 and fibrillarin are normally present within wild type MB neuroblasts relative to other neuroblasts. As an initial query, we analyzed the NB-lineage specific transcriptome data set from Yang et al. (2016) who performed a cell-type specific RNA-seq analysis for *Drosophila* larval neuroblasts (non-selective “all” NBs, MB NBs, AL NBs, and Type II NBs), neurons, and glia. We found the expression levels of four RBFs (*Nopp140, fibrillarin, Nop56*, and *Nop60B*) were higher in the MB NBs compared to the AL NBs and Type II NBs (Fig. 9A). We also checked the expression levels of the four *Drosophila* Nucleostemin orthologues (NS1 – NS4) (Kaplan et al., 2008; Rosby et al., 2009; Hartl et al., 2013; Wang and DiMario, 2017). Mammalian nucleostemin (NS) is a nucleolar GTP-binding protein originally described in embryonic and neuronal stem cells and in certain cancer cells that can regulate both ribosome production and cell cycle progression (Tsai, 2011). We found *NS1, NS2*, and *NS4*, but not *NS3* expressed at higher levels in the neuroblasts than in neurons (Fig. 9B). As controls, we checked the expression levels of *Deadpan* which encodes a NB-specific transcription factor, *Prospero* which encodes a NB- and GMC-specific transcription factor, and *Elav* which encodes a *Drosophila* neuron-specific protein. As expected, *Deadpan* and *Prospero* expression levels were higher in NBs compared to neurons, and *Elav* expression was higher in neurons compared to NBs (Fig. 9C).

**Fig. 9.**
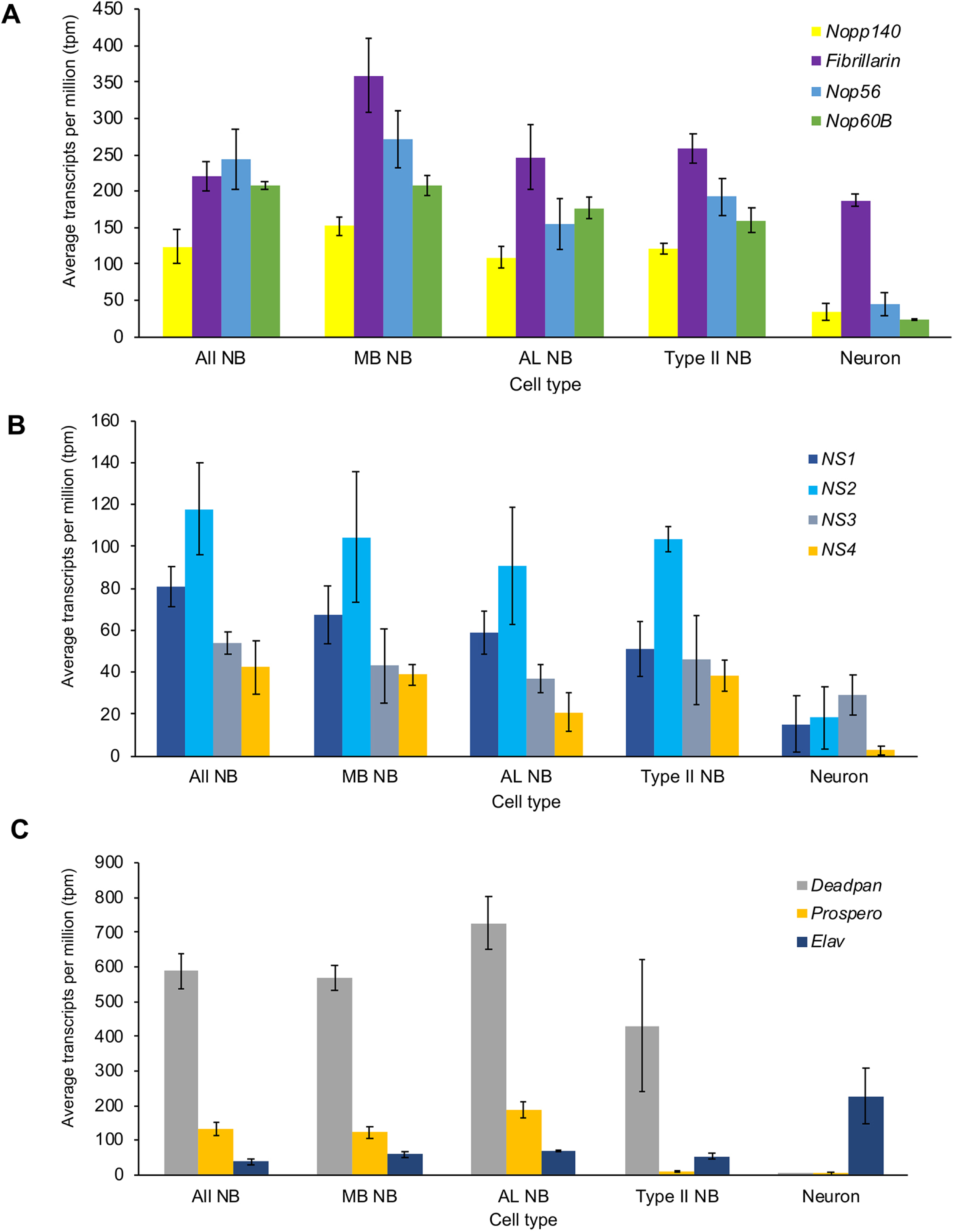
Transcriptome analyses of ribosome biogenesis factors in lineage-specific *Drosophila* neuroblasts and neurons. Expression levels of *Nopp140, fibrillarin, Nop56*, and *Nop60B* transcripts (A); *NS1, NS2, NS3*, and *NS4* transcripts (B); *Deadpan, Prospero*, and *Elav* (C) in the *Drosophila* larval NBs and neurons. All NB (n=3), Mushroom Body (MB) NB (n=3), Antennal Lobe (AL) NB (n=3), Type II NB (n=3), neurons (n=2). Transcriptome data obtained from Yang et al. (2016).

To inquire if differences exist in the expression levels for genes encoding ribosomal proteins in different NB lineages, we again analyzed the transcriptomic data set of Yang et al. (2016). Out of 92 genes encoding ribosomal proteins (recall that ~80 different proteins constitute an intact ribosome), 47 genes were preferentially expressed in MB NBs compared to AL NBs, Type II NBs, and neurons (Supplemental Fig. S5). Noticeably, *RpL41* had the highest expression levels among all ribosomal protein genes in all brain cell types examined, but *RpL41* transcripts levels were significantly higher in the MB NBs (indicated by an asterisk in Supplemental Fig. S5). Enhanced expression of *RpL41* and the other 46 genes in MB NBs suggests that their ribosomes may be different from those in other NB lineages.

## DISCUSSION

A wealth of knowledge exists for *Drosophila* neurogenesis making it possible to analyze developing brains at the level of individual neuroblast lineages (Birkholz et al., 2015; Egger et al., 2008; Hartenstein and Wodarz, 2013; Homem and Knoblich, 2012; Urbach and Technau, 2003). We asked if Type I and II neuroblasts, Mushroom Body (MB) neuroblasts, Optic Lobe neuroblasts, and Antennal Lode (AL) neuroblasts are affected variably upon nucleolar stress, as are stem cells and precursor cells in the human ribosomopathies. To induce nucleolar stress, we used CRISPR-Cas9 to disrupt the *Nopp140* gene which encodes two isoforms that function as early RBFs. *Drosophila* larvae homozygous for the *Nopp140* disruption allele, *J11^non-DsRed^*, showed smaller brains by day 4 ALH (Fig. 5A). These *Nopp140−/−* larvae arrested in the second instar stage, and while some lingered to day 24 ALH; none of them survived (Fig. 4B). Compared to wild-type brains, *Nopp140−/−* larval brains at day 2-3 ALH had far fewer proliferating NBs. However, deadpan antibody labeling and EdU labeling of homozygous *J11^non-DsRed^* larvae showed that MB neuroblasts, and in some cases the AL neuroblasts, proliferated through the embryo-to-larva transition and continued to proliferate at day 2-3 ALH as other neuroblast lineages remained arrested (Fig. 6). Hence, MB NBs exhibited resilience to nucleolar stress due to loss of zygotic *Nopp140* gene expression.

### Ontogenesis of the Mushroom Body Neuroblast Lineages

Insect MBs are central hubs for olfactory sensory input, learning, and memory (Thum and Gerber, 2019). Formation of the MBs begins during embryogenesis during which each MB NB differentially expresses unique combinations of the regulatory genes (Kunz et al., 2012; Yang et al., 2016). As far as we know, none of these gene products have direct links to ribosome production. During the embryo-to-larva transition, only the four MB NBs and one Antennal lobe (AL) NBs continue to proliferate independently of dietary nutrients and PI3-kinase activity (Kunz et al., 2012; Prokop and Technau, 1994; Ito and Hotta, 1992; Lin et al., 2013; Sipe and Siegrist, 2017). The majority of other NBs, however, enter a period of quiescence (Kunz et al., 2012; Prokop and Technau, 1994; Ito and Hotta, 1992) and require dietary nutrients and PI3-kinase activity to exit quiescence ~24 hrs after hatching (Lovick and Hartenstein, 2015; Sipe and Siegrist, 2017). During subsequent larval development, each MB NBs generates an almost identical repertoire of intrinsic Kenyon cells and continues to proliferate on into the pupal stages (~85-90 hours after pupa formation) (Ito and Hotta, 1992; Ito et al., 1997). As a result, the adult MB neuropil in each CB lobe is densely packed with around 2000-3000 Kenyon cells per lobe (Technau and Heisenberg, 1982; Aso et al., 2009).

Thus the MB NBs (and AL NBs) are inherently different in their proliferative schedules compared to the rest of the neuroblasts in the *Drosophila* larval brain. This may explain in part their resilience to nucleolar stress; that is, continued neuroblast proliferation and already high synthetic rates (e.g., ribosome production) may sustain the MB NBs upon nucleolar stress at least temporarily, while the other NBs may not be able to rekindle high synthetic levels as they exit quiescence (Bertoli et al., 2013).

### Phenotypes

For the first time, we showed that the *Nopp140* gene in *Drosophila* is haplo-insufficient where *J11^DsRed^/TM3* displayed embryonic lethality (Fig. 4A). This was a surprise since a previous segmental aneuploidy study indicated no haplo-insufficiency genes existed in cytological region 78F4 of the left arm of chromosome 3 (Lindsley et al., 1972). Haplo-insufficiency of the *Drosophila Nopp140* gene would be analogous to haplo-insufficiency of the human *Tcof1* gene which encodes treacle, a vertebrate early RBF related to Nopp140 in structure and function. Loss of treacle in *Tcof1+/−* human embryos results in the Treacher Collins syndrome, a ribosomopathy leading to apoptosis in select embryonic neural crest cells ultimately leading to the craniofacial birth defects.

Earlier work in our lab showed that complete loss of Nopp140 in *Drosophila* induced nucleolar stress with the redistribution of the C/D box methyl-transferase, fibrillarin, to the nucleoplasm (He et al., 2015). Here, we showed that *Nopp140* transcripts and at least the Nopp140-RGG isoform were reduced but not completely absent in early larvae homozygous for the disrupted *Nopp140* allele, *J11^non-DsRed^*, (Figs 2, 3). Interestingly, each wild type central brain lobe showed four anterior cells that appeared to contain more Nopp140-RGG than other cells within the same lobes (Fig. 3). The observation suggested that MB NBs contain more Nopp140 than do other NBs. We then showed that mCD8::GFP-labeled MB lineages in homozygous *J11^non-DsRed^* larvae retained nucleolar fibrillarin, whereas fibrillarin was noticeably redistributed to the nucleoplasm in the majority of other NBs (Fig. 8). This latter observation indicated that nucleoli in the MB lineages preferentially retained more RBFs and perhaps maintained functional production of ribosomes longer, although to what extent requires future molecular analyses.

### Differential Expressions

Most cells within the central brain lobes of homozygous *J11^non-DsRed^* larvae 1-2 day ALH showed reduced anti-Nopp140-RGG labeling compared to brain cells in similarly aged wild type larvae. Interestingly, wild type larvae clearly showed four cells, identified as MB NBs, per central brain lobe with prominent anti-Nopp140-RGG labeling (Fig. 3). The observation suggests that wild type MB NBs contain more zygotically expressed Nopp140 than do other NBs. Recent findings supporting this notion show that various RBFs such as treacle, fibrillarin, Nop56, mbm, and NS3 are overexpressed in stem cell and progenitor cell populations (Brombin et al., 2015; Watanabe-Susaki et al., 2014; Wang et al., 2013; Johnson et al., 2018; Hovhanyan et al., 2014; Hartl et al., 2013; Dixon et al., 2006), and that the quantity and spatiotemporal expression of RBFs can vary in different stem cell or progenitor cell populations (Weiner et al., 2012; Bouffard et al., 2018). A selective expression of not only Nopp140, but other RBFs such as fibrillarin, Nop56, Nop60b (Fig. 9A) in MB NBs during the embryo-to-larva transition period may explain the resilience of MB lineages to nucleolar stress later during larval development.

Related to differential expression of RBFs in stem cells and progenitor cells is the perplexing problem of why some larvae homozygous for the disrupted *Nopp140* gene survive up to day 24 ALH (Fig. 4B). While we have yet to pursue this question rigorously, we suspect these *Nopp140−/−* individuals may inherent more maternal *Nopp140* mRNA and/or protein, and this may be a function of the nutrition and health of the mothers. Earlier work in our lab (McCain et al., 2006) followed maternally expression of GFP-Nopp140 into embryogenesis, and we noted then that the protein has a long lifespan, on the order of several days. Individual *Nopp140−/−* embryos that inherited extra maternal *Nopp140* transcripts or protein would likely produce more ribosomes and survive longer.

### The Possibility of Heterogeneous Ribosomes

Ribosomes are not all the same even within a single cell (Xue and Barna, 2012; Guo 2018). Werner et al. (2015) showed that the translation program of human embryonic stem cells (hESCs) differentiating into neural crest cells changed after the depletion of KBTBD8, a substrate adapter for the vertebrate-specific ubiquitin ligase, CUL3. CUL3 mono-ubiquitylates human Nopp140 (NOLC1) and treacle, and forms a Nopp140-treacle platform that connects RNA Pol I machinery with ribosome modification factors. Based on these results, the authors hypothesized that the change in translational profile was the result of differential alteration of ribosomes. Modifications such as rRNA pseudouridylation and methylation, or phosphorylation and ubiquitylation of ribosomal proteins or ribosome-associated factors may ultimately contribute to translational control of gene expression (Sloan et al., 2017). Thus, the abundance of Nopp140 in different *Drosophila* neuroblasts could potentially lead to differential modifications of the ribosome pool, and thus changes in the translational profile in different neuroblasts.

Finally, transcriptome profiles (Fig. 9 and Supplemental Fig. 5) of the different *Drosophila* larval brain neuroblasts indicated that RBFs fibrillarin, Nop140, Nop60, and Nop56 are expressed at higher levels in the MB NBs. Of the ribosomal proteins, RpL41 had the highest expression levels in all brain cells, but was again highest in the MB NBs (indicated by an asterisk in Supplemental Fig. S5). Are there heterogeneous pools of ribosomes within a *Drosophila* larval brain? If so, it might explain why the MB NBs are more resilient to nucleolar stress. The *Drosophila* nervous system should serve well to explore differential threshold requirements for ribosome production.

## Competing Interests

The authors declare no competing or financial interests.

## Funding

This study was funded by the National Science Foundation, grant numbers MCB0919709 and MCB1712975.

